# Characterization of *Dnajc12* knock-out mice, a model of hypodopaminergia

**DOI:** 10.1101/2024.07.06.602343

**Authors:** Isaac Bul Deng, Jordan Follett, Jesse D. Fox, Shannon Wall, Matthew J. Farrer

## Abstract

Homozygous *DNAJC12* c.79-2A>G (p. V27Wfs*14) loss-of-function mutations were first reported as a cause of young-onset Parkinson’s disease. However, bi-allelic autosomal recessive pathogenic variants in *DNAJC12* may lead to an alternative constellation of neurological features, including infantile dystonia, developmental delay, intellectual disability and neuropsychiatric disorders. DNAJC12 is understood to co-chaperone aromatic amino acid hydroxylases to foster the synthesis of biogenic amines. *In vitro*, we discover overexpressed DNAJC12 forms a complex with guanine triphosphate cyclohydrolase 1 (GCH1), the rate-limiting enzyme in the synthesis of tetrahydrobiopterin, a cofactor paramount for biogenic amines synthesis. We also confirm DNAJC12’s interaction with tyrosine (TH) and tryptophan hydroxylase (TPH), which are rate-limiting enzymes for synthesis of biogenic amines dopamine (DA) and serotonin (5-HT). *In-vitro* knock-down of DNAJC12 with a siRNA destabilizes the DNAJC12-TH-GCH1 complex, reducing GCH1 levels, whereas reciprocal overexpression of both TH and GCH1 increases endogenous DNAJC12, alluding to the significance of modulating the DNAJC12-TH-GCH1 complex as a therapy for DNAJC12 and other biogenic amine disorders. We extend these investigations to a Cre-conditional knock-out mice (cDKO) in which loxP sites flanking *Dnajc12* exon 2 enable its excision by cre-recombinase. With germline Cre expression, we have created a constitutive *Dnajc12* knock-out (DKO). DKO mice exhibit reduced locomotion/ exploratory behavior at 3 months in automated open-field testing, accompanied by increased plasma phenylalanine which is a cardinal feature of patients with pathogenic *DNAJC12* variants. In striatal tissue, total DA and 5-HT, their metabolites, and electrically-evoked DA release are all reduced. Biochemical alterations in synaptic proteins are also apparent, with enhanced phosphorylation of Th pSer31 and pSer40 reflecting biological compensation. Most immediately, cDKO and DKO mice present models to develop and refine therapeutic approaches for biogenic amines disorders, including dystonia and parkinsonism. They will also enable the pleiotropic functions of biogenic amines (including DA), usually synthesized in the brain or periphery, to be separated.

## Introduction

A major pathological hallmark of neurodegenerative diseases is the presence of intracellular and extracellular protein inclusions, indicative of altered proteostasis.^1,2^ The maintenance of cellular proteome requires molecular chaperones known as heat shock proteins (HSP), which comprise of HSP90, HSP70, HSP60, HSP40 and small HSPs. This diverse superfamily has a fundamental role in protein folding, trafficking and degradation.^1, 2^ Biallelic, recessively inherited pathogenic variants in *DNAJC12*, a member of the *DNAJC* family, also referred to as HSP40, induce a multitude of neurological disorders, including young-onset parkinsonism, infantile dystonia, developmental delay, intellectual disability and neuropsychiatric disorders.^3–9^ Variants in *DNAJC12* also account for a small proportion (∼5%) of cases of hyperphenylalaninemia (HPA), which is prominently attributed to variants in *phenylalanine hydroxylase* (*PAH*), and its cofactor *tetrahydrobiopterin* (*BH_4_*).^4,5,9,10^Although its function has yet to be fully elucidated, DNAJC12 is widely regarded as a co-chaperone for aromatic amino acid hydroxylases (AAAHs) including PAH, tyrosine hydroxylase (TH) and tryptophan hydroxylase 1 and 2 (TPH1 and TPH2).^1,3,4^ Patients with *DNAJC12* variants have reduced levels of dopamine (DA) and serotonin (5-HT) metabolites in their cerebrospinal fluid, congruent with the role of DNAJC12 as a co-chaperone for TH and TPH2, respectively.^3–6,9^

Several members of the DNAJC family of co-chaperones besides DNAJC12 are implicated in parkinsonism, including cysteine string protein-α (DNAJC5), auxilin (DNAJC6), receptor-mediated endocytosis-8 (DNAJC13) and cyclic G associated kinase (DNAJC26).^11–14^ We have previously identified pathogenic variants in *guanosine triphosphate (GTP) cyclohydrolase* I (*GCH1*), the rate-limiting step in the conversion of GTP to tetrahydrobiopterin (BH_4_). Heterozygous patients with mutant *GCH1* either present with postural dystonia in childhood that becomes generalized^15,16^ or with late-onset Parkinson’s disease (PD).^17,18^ The former appears to be a *forme fruste* of the latter; moreover, as both occur within different individuals of the same families, with the same mutations, within genes in intersecting pathways. What underlies this profound difference in penetrance and presentation remains enigmatic.^19^ Dopaminergic neurons in the *substantia nigra par compacta* (*SNpc,* located in MB) extend their axons via the medial forebrain bundle to the striatum (STR), where DA is released as a volumetric neuromodulator of synaptic transmission.^20^Clearly, dystonia and/or parkinsonism in patients with *DNAJC12*^3–5,21,22^ or *GCH1*^15,17,23,24^ pathogenic variants can be attributed to DA dysregulation as each is responsive to levodopa (L-DOPA) treatment, a DA-replacement therapy. To understand the mechanistic role of DNAJC12 in DA synthesis requires *in-vivo Dnajc12* models, as further study is likely to reveal novel strategies to re-establish DA levels to help alleviate dystonia and parkinsonism. The first homozygous pathogenic variant discovered in human disease was *DNAJC12* c.79-2A>G (p. V27Wfs*14), which results in exon 2 skipping, transcript frameshift and nonsense-mediated decay.^5^ Normally, exons 2 and 3 encode the DNAJC and Hsp70 interaction domains. Here we describe the generation and first characterization of murine *Dnajc12* knock-out mice (Dnjac12 p. V27Wfs*44) focused on the role of Dnajc12 in DA synthesis, physiology and related behavior.

## Methods

### Animals

The gene locus for murine *Dnajc12* is on chromosome 10 at 63,218,222-63,244,619 (GRCm39: GCF_000001635.27). The main protein encoding isoform 1 encodes 198 amino acids, denoted by NCBI transcript RefSeq ID NM_013888.3. It comprises 5 exons and generates a mRNA of 1305bp. There is a second shorter isoform 2 (NM_001253685.1) that uses a different exon 1, encoding 116 amino acids and lacking a DNAJC domain. As in humans, deletion of exon 2 in the murine sequence (ENSMUSE00001301217, Chr10:63,231,564-63,231,642 [GRCm39]) is also predicted to alter the open reading frame after amino acid 26 (exon 1) and result in 17 novel amino acids that terminate in a stop mutation (p. V27Wfs*44). Hence, using a C57BL/6J background, Jackson Laboratories used CRISPR/Cas9 to insert a pair of loxP sites flanking exon 2 to create a conditional allele and mouse strain (cDKO) (Fig S1). Germline Cre recombination was subsequently used to ablate the intervening sequence and create a constitutive knock-out (DKO). All subsequent animal procedures at the University of Florida were performed in accordance with a protocol approved by the Institutional Animal Care and Use Committee. DKO mice were maintained by heterozygous breeding on a reverse light cycle (8 pm to 8 am) and single-sex group housed in enrichment cages after weaning at post-natal day 21. All mice were supplied with a standard rodent chow and water *ad libitum*. Ear notches were taken at *post-mortem* for *post-hoc* genotyping with the following primer sequences 5’-CCAGGCCTGGCTTTTATTTG-3’ and 5’-TTAAGTCCAAACCAGCTGCC-3’ using standard thermocycling conditions. All experiments, data processing and analysis were conducted in a blinded manner.

### Behavior

#### Open field test

The Activity Monitor System uses an array of infrared (IR) photo beams to measure locomotion and motor activity in a square open field arena with a duration of 30 min. The apparatus consists of 3 sets of matching IR photo beams that project across the open field along x, y and z axes. The software detects when and where the animal disrupts the photo beams, and allows the open field arena to be broken into areas of interest depending on the experiment.^25^

#### Grip Strength test

Neuromuscular function and muscular strength were assessed by measuring the peak force required to make the animal release its grip.^26^ The Grip Strength test apparatus consists of a baseplate, a trapezoidal stainless-steel grip, and a force sensor. To measure grip strength, mice are gently lifted by the tail so that the forelimbs or hindlimbs can grasp onto the steel grip and then gently pull away from the apparatus by the tail until they release their grip.

#### Balance beam

Sensorineural balance and coordination were examined by placing mice on an elevated narrow square beam of 100 cm in length, with an enclosed dark platform at the end of the beam.^27,28^ Mice are trained on a beam with a width of 24 mm and performance is tested at 12 mm. Videos are recorded from both sides of the beam and total foot slips are measured.

#### Antibodies

Rabbit monoclonal or polyclonal antibodies (supplier, dilution) include those against: 14-3-3 (Invitrogen, 51-0700, 1:1000); clathrin heavy chain (Abcam, ab21679, 1:1000); DNAJC5 (Synaptic systems, 154003, 1:1000); DNAJC12 (12338-1-AP, Proteintech; 1:1000); Endophilin 1 (Synaptic systems, 159002, 1:2000); GFP (Abcam, ab290; 1,10,000); HSC70 (Abcam, ab51052; 1:1000); myelin basic protein (Abcam, ab218011, 1:1000); olig2 (Abcam, ab109186, 1:1000); Synaptobrevin 2 (VAMP2) (Synaptic systems, 104202, 1:1000); tyrosine hydroxylase (pser40) (Sigma, T9573, 1:1000); tyrosine hydroxylase (pSer31) (Sigma, SAB4300674, 1:1000); β-tubulin (Abcam, ab6046; 1:4000). Mouse monoclonal antibodies include those raised against: GAPDH (ThermoFisher Scientific, MA5-15738; 1:5000), FLAG (M2 clone, Millipore-Sigma, F3165; 1:5000); tyrosine hydroxylase (TH) (Sigma, T2928, 1:1000); SNAP25 (Synaptic systems, 111011, 1:1000). Rat monoclonal antibodies: MYC (Abcam, ab206486)

#### Tissue collection and homogenization

Mice were deeply anaesthetized via inhalation of isoflurane and then intracardially perfused with phosphate buffer saline (PBS) prior to microdissection of the MB and STR. For analysis of total protein, micro-dissected tissues were homogenized at 20% w/v in 1x Cell Signaling Lysis Buffer (#9803S) with 1x Halt™ Protease and Phosphatase Inhibitor Cocktail (Thermofisher #78442) using a 1 mL glass Dounce homogenizer (Wheaton) and placed on ice for 30 min before being cleared by centrifugation (14,000 ×g at 4°C for 20 min). The total protein concentration in the supernatant was measured with BCA assay (Pierce).^29,30^

#### Cell culture

HEK293FT (Thermofisher Scientific; R70007) cell line was maintained in high glucose Dulbecco’s Modified Eagles Medium (DMEM), respectively, supplemented with 10% (v/v) FBS (Gibco), 2 mM l-glutamine (Thermo Fisher Scientific) and penicillin/streptomycin. Cultures were maintained in 5% CO_2_ at 37°C.^31^ Transient transfection of cDNA was performed using Lipofectamine 2000 (Thermo Fisher Scientific), while siRNA was transfected using Lipofectamine RNAiMAX (Thermofisher Scientific) according to the manufacturer’s instruction.

#### Co-immunoprecipitation

For experiments where epitope-tagged constructs were transiently overexpressed, HEK293 cells were seeded in 10 cm dishes and transfected with the indicated plasmids (GFP-DNAJC12, FLAG-TH, FLAG-TPH1 and MYC-GCH1). At 48h post-transfection, cells were washed once in ice-cold PBS and lysed in 1 mL of Cell lysis buffer (Cell signaling #9803S), supplemented with 1x Halt™ Protease and Phosphatase Inhibitor Cocktail (Thermofisher #78442) and incubated on ice for 30 min with gentle agitation every 15 min before being cleared by centrifugation at 14,000 ×g for 12 min at 4°C. 20 μL of cleared lysate was collected as input, and GFP was recovered from the lysate using 20 μL of pre-equilibrated GFP-Trap (gta-20, ChromoTek) for 1h at 4°C. Beads were collected by centrifugation and washed 4-6 times with 1 mL of lysis buffer followed by centrifugation before elution (10 min, 90 °C) in 2 x SDS-loading buffer prior to standard SDS-PAGE and western blotting.

#### SDS-PAGE and Western blotting (WB)

20-40 µg of denatured lysates were resolved by SDS-PAGE on 4-20% TGX Criterion TGX pre-cast gels (Bio-rad) and transferred to nitrocellulose membranes by semi-dry trans-Blot Turbo transfer system (Bio-rad). The membranes were blocked with either 50% Intercept tris buffered saline (TBS; 927-60001), 5% bovine serum albumin (BSA) in TBS-Tween (TBST), or 5% milk in TBST for 1h at room temperature (RT) followed by incubation overnight at 4°C with primary antibodies diluted in blocking buffers depending on the protein of interests. After incubation with primary antibodies, the membranes were washed in TBST (3 x 5 min), followed by incubation for 1h at RT with fluorescently conjugated goat anti-mouse or rabbit IR Dye 680 or 800 antibodies (Licor). The blots were washed in TBST (3 x 5 min) and scanned on a ChemiDoc MP imaging platform (Bio-Rad).^29,30^

#### Immunofluorescence with confocal microscopy

For imaging, mice were anesthetized via inhalation of isoflurane, intracardially perfused with PBS and their brains were rapidly extracted then post-fixed overnight at 4°C in 4% paraformaldehyde (PFA). Cryoprotection of the brains was performed with an increasing sucrose gradient (10%, 20%, 30% in PBS) before sagittal slices (30 µm) were obtained with a cryostat. Sections were blocked in 10% normal goat serum (NGS) in 0.3% PBST (1h, 37°C), followed by incubation of primary antibodies: rabbit anti-DNAJC12 (12338-1-AP, Proteintech; 1:200), chicken anti-Th (ab76442 Abcam; 1:500) and guinea pig anti-Cre (1:250, Cell signaling) made in antibody solution (5% NGS + 0.3% PBST) overnight at 4°C. The sections were washed (4×15 min in PBS), followed by secondary antibody incubation with species-specific Alexafluor IgG secondary antibodies (1h at RT, Invitrogen; 1:500). Tissues were rewashed in PBS (4 × 15 min), then co-stained with DAPI (ThermoFisher Scientific,1:5000) and mounted using Fluoromount-G® (0100-20, Southern Biotech). Images were acquired using a 20x objective (Cre) or a 60x oil objective on an Olympus Fluoview FV-1000 confocal laser scanning microscope (9 x 0.33μm step size) (Dnajc12). Images were stacked (three stacks of 3 individual images viewing 1 μm tissue depth) using FIJI ImageJ software (NIH, USA).^29,30^

#### Electrophysiology

Fast scan cyclic voltammetry (FSCV) was used to assess evoked DA release and reuptake in 300 μm hemi-coronal slices of the dorsolateral striatum, as previous.^29,30^ Coronal striatal slices obtained from 3M DKO mice with a vibratome were perfused with oxygenated artificial cerebrospinal fluid (aCSF) at RT for ∼1h prior to the experiments. Stimuli are delivered by nickel-chromium bipolar electrodes (made in house) placed in the dorsolateral striatum, optically isolated (World Precision Instruments, USA) and controlled/sequenced with ClampEx software. Voltammetric responses were recorded, standardized and analyzed with an Invilog Voltammetry system and software (Invilog), using carbon fiber electrodes (diameter: 32μm, length: 30μm, sensitivity: 21-40nA/μM; Invilog) and placed within 100-200 μm of the stimulating electrode in the dorsolateral striatum. Field activity was recorded to ensure the viability of the slice during the recording. Input/output paradigms were as previous.^29,32^ At the end of each recording session, a three-point calibration of each carbon fiber electrode is conducted (final concentrations of DA in a CSF (μM): 0.5,1.0 and 2).

#### High-performance liquid chromatography (HPLC)

Sample preparation and HPLC were performed by the Emory HPLC Bioanalytical Core (EHBC) according to their protocol. In brief, striatal tissues were suspended in 300 μL solution of 0.1M PCA and 0.1mM EDTA, followed by sonication, then centrifugation of the homogenates at 10,000 xg for 15 min at 4°C to obtain the supernatant. The concentration of DA,5-HT and their metabolites in the supernatant were determined by reverse phase HPLC with electrochemical detection using an ESA 5600A CoulArray detection system equipped with an ESA Model 584 pump and an ESA 542 refrigerated autosampler.

#### Metabolomics

Plasma samples for DKO and WT mice were sent to Creative Proteomics to analyze free amino acids and related metabolites, as per their protocol. To summarize, 20 μL of each plasma sample was added into an Eppendorf tube containing 80 μL of the internal standard solution (contains 41 isotope-labeled internal standards). The resultant mixture was sonicated for 5 min and centrifuged at 21,000 xg for 10 min before taking the supernatant. 50 μL of supernatant for each sample or the standard calibration solution was mixed with 100 μL of 20 mM dansyl chloride solution and 100 μL of borate buffer (pH 9), and allowed to react for 30 min at 40°C. Aliquots of the resultant solutions (5 μL) were injected to run UPLC-MRM/MS on a Waters Acquity UPLC system coupled to a Sciex QTRAP 6500 Plus mass spectrometer equipped with a heated ESI source. LC separation was carried out on a C18 UPLC column (2.1*150 mm, 1.8 μm) with 0.1% formic acid in water and 0.1% formic acid in an acetonitrile-isopropanol mixed solvent as the binary-solvent mobile phase for gradient elution (20% to 90% B in 20 min), at 55 °C and 0.3 mL/min.

#### Statistical analysis

The data for the automated open field test was acquired with Activity Monitor 7 software (Med Associates Inc). Data acquisition and extraction of the spike corresponding to the force exerted by mice was analyzed with Grip Strength Software (Maze Engineers). DA traces for FSCV were analyzed with Clampfit (Molecular devices), and the densitometric analyses were performed with Image J software. Normality testing and statistical analysis of all the data were performed using GraphPad Prism 10. The data is presented as mean ± standard error of the mean (SEM).

## Results

### DNAJC12 interacts with TH, TPH, GCH1 and HSC70 *in-vitro*

Recent unbiased mass-spectrometry in HEK293 cells overexpressing the co-chaperone DNAJC12 revealed its interaction with DA synthesizing enzyme tyrosine hydroxylase (murine ‘Th’, human ‘TH’).^4^ To build upon this data, HEK293 cells transiently co-transfected with combinations of epitope-tagged *DNAJC12* (GFP-DNAJC12)*, TH* (FLAG-TH), TPH1 (FLAG-TPH1) and GCH1 (MYC-GCH1) were lysed and immunoprecipitated with GFP-agarose beads. GFP-tagged DNAJC12 co-immunoprecipitated FLAG-TH (Fig 1a,d), FLAG-TPH1 (Fig 1b,c,e), MYC-GCH1 (Fig 1c) and endogenous HSC70 (Fig 1a,b, c, d and e), while empty vector GFP control did not, consistent with recent mass-spectrometry findings.^4^ Knock-down of DNAJC12 in HEK293 with a siRNA destabilized the DNAJC12-TH-GCH1 complex, reducing GCH1 levels in the process, while the amount of TH was unaltered (Fig 1f-g). Unexpectedly, co-overexpression of TH and GCH1 increased levels of endogenous DNAJC12, while their individual expression had no effects (Fig 1h). To confirm this interaction is not limited to overexpression systems, lysates generated from sub-cortical brain tissue of wildtype mice (WT) were immunoprecipitated with anti-Th antibody-coated beads, then eluted and subjected to WB analysis, and gave complementary findings as in Fig S1a. Fluorescence immunohistochemistry (IHC) with confocal microscopy also visualized the sub-cellular localization of Dnajc12 within dopaminergic neurons (DA-producing neurons, identified with Th antibody) in MB slices from WT mice. Dnajc12 fluorescent puncta were observed within the soma, dendrites and axons of Th-positive dopaminergic neurons in the MB (Fig S1b).

**Fig 1.**
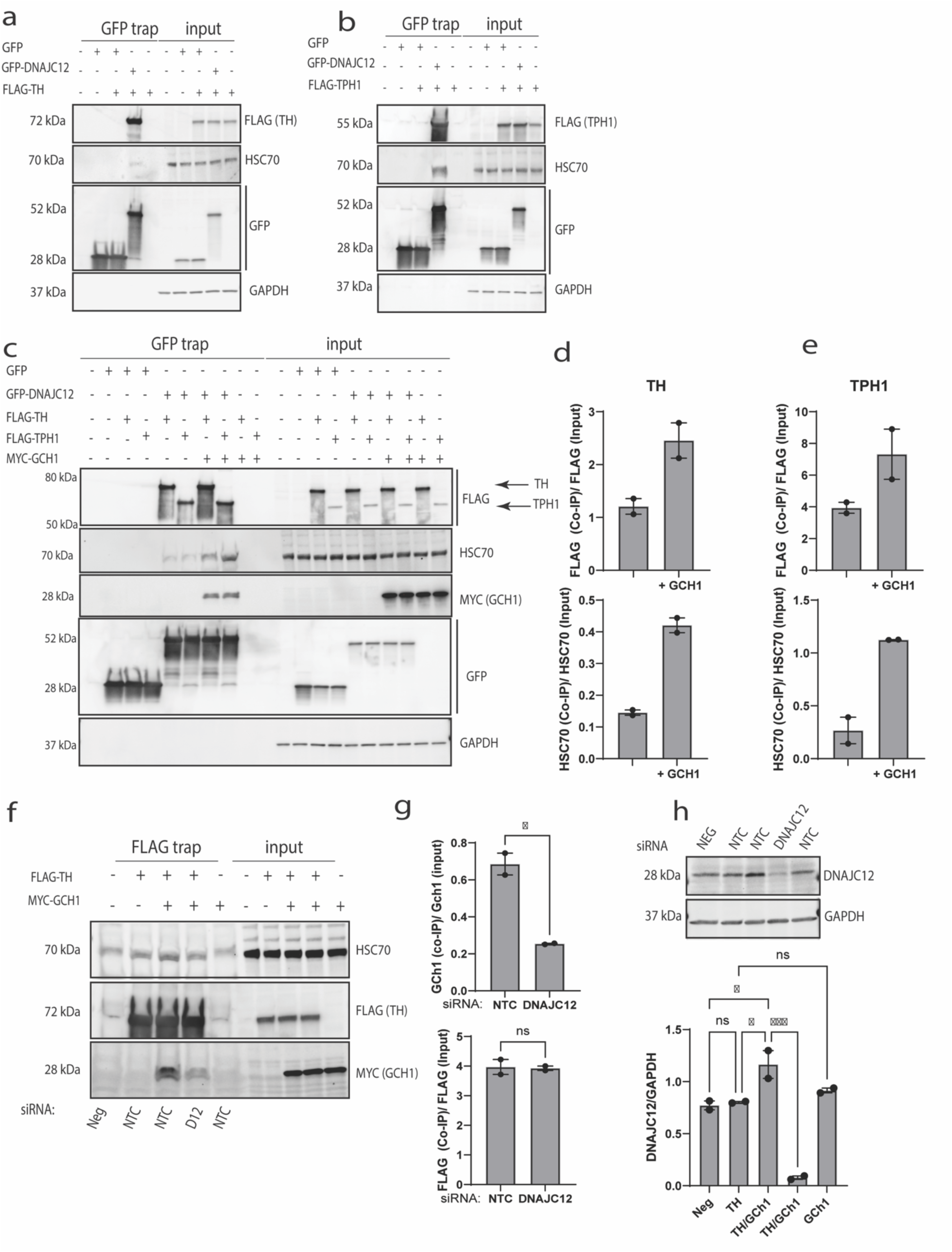
DNAJC12 co-immunoprecipitated TH, TPH1, GCH1 and Hsc70 in HEK293FT cells. HEK293 cells transiently co-transfected with combinations of epitope-tagged *DNAJC12* (GFP-DNAJC12)*, TH* (FLAG-TH), TPH1 (FLAG-TPH1) and GCH1 (MYC-GCH1) were lysed and immunoprecipitated with GFP-agarose beads. The products were then resolved with SDS-page and probed with the following antibodies. GFP-DNAJC12 immunoprecipitated FLAG-TH (**a**), FLAG-TPH1 (**b**) and endogenous Hsc70 (**a, b**). Representative blot (**c**) and quantification of GFP-DNAJC12 co-immunoprecipitation of TH with Hsc70 (**d**), and TPH with Hsc70 (**e**) when MYC-GCH1 is overexpressed (**g**). Representative blot (**f**) and FLAG-DNAJC12 co-immunoprecipitation of MYC-GCH1 and FLAG-TH when DNAJC12 is knock-down with SiRNA (**g**). Quantification of endogenous Dnajc12 when TH and GCH1 are overexpressed (**h**).

### DKO mice have impaired locomotion and exploratory behavior in open field testing at 3M

We investigated the effects of Dnajc12 ablation (Fig 2a) on nigrostriatal DA synthesis, physiology and behavior, given individuals with pathogenic *DNAJC12* variants respond well to DA-replacement therapy. We confirmed the ablation of *Dnajc12* with genomic PCR (Fig 2b) and loss of protein expression in MB (where the cell bodies of dopaminergic neurons reside) and STR (location of their terminals) (STR) (Fig 2c-e). We then characterized the behavior of DKO mice at 3M relative to WT littermates using an automated open-field test. DKO mice had a significant reduction in their ambulatory distance (Fig 3b), ambulatory time (Fig 3c), number of rears (Fig 3e) and average speed (Fig 3f), along with an increase in resting time (Fig 3d), all of which are indicative of impaired locomotion/exploratory behavior. There were no deficits in grip strength (Fig 3g) or motor coordination on a beam (Fig 3h-i).

**Fig 2.**
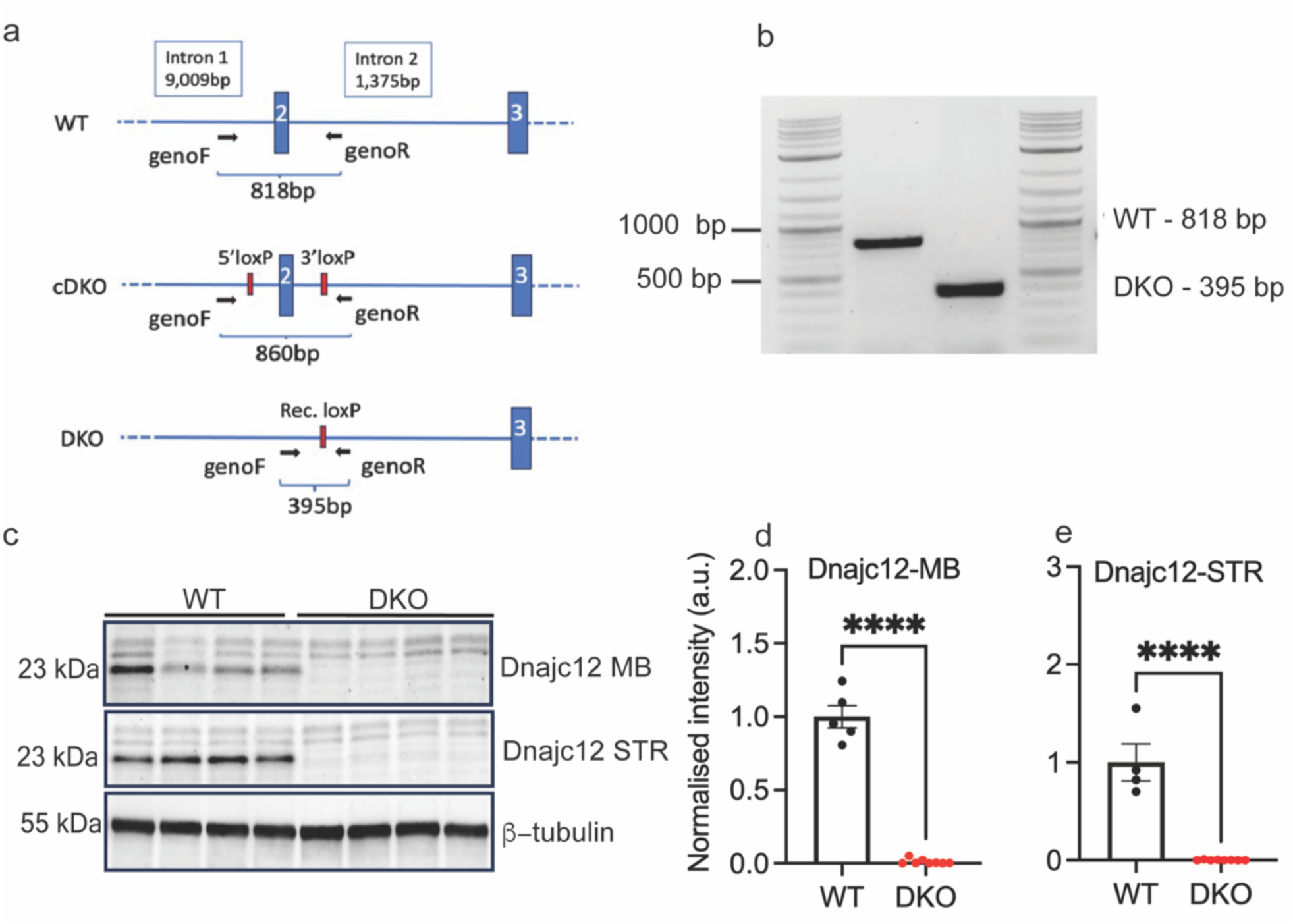
Depiction of *Dnajc12* ablation in 3M DKO mice. Illustration of cre-mediated excision of *Dnajc12 (**a**).* Genomic PCR for *Dnajc12* (**b**). Representative blots for Dnajc12 expression in the midbrain (MB) and striatum (STR) (**c**). Densitometric analyses of Dnajc12 expression in the MB (**d**, unpaired *t*-test, t_11_ = 16.69, p <0.0001) and striatum (**e**, unpaired *t*-test, t_10_ = 7.80, p <0.0001). *\*\*\*\*p < 0.0001.* Error bars are ± SEM. WT n =4-5, DKO n = 8.

**Fig 3.**
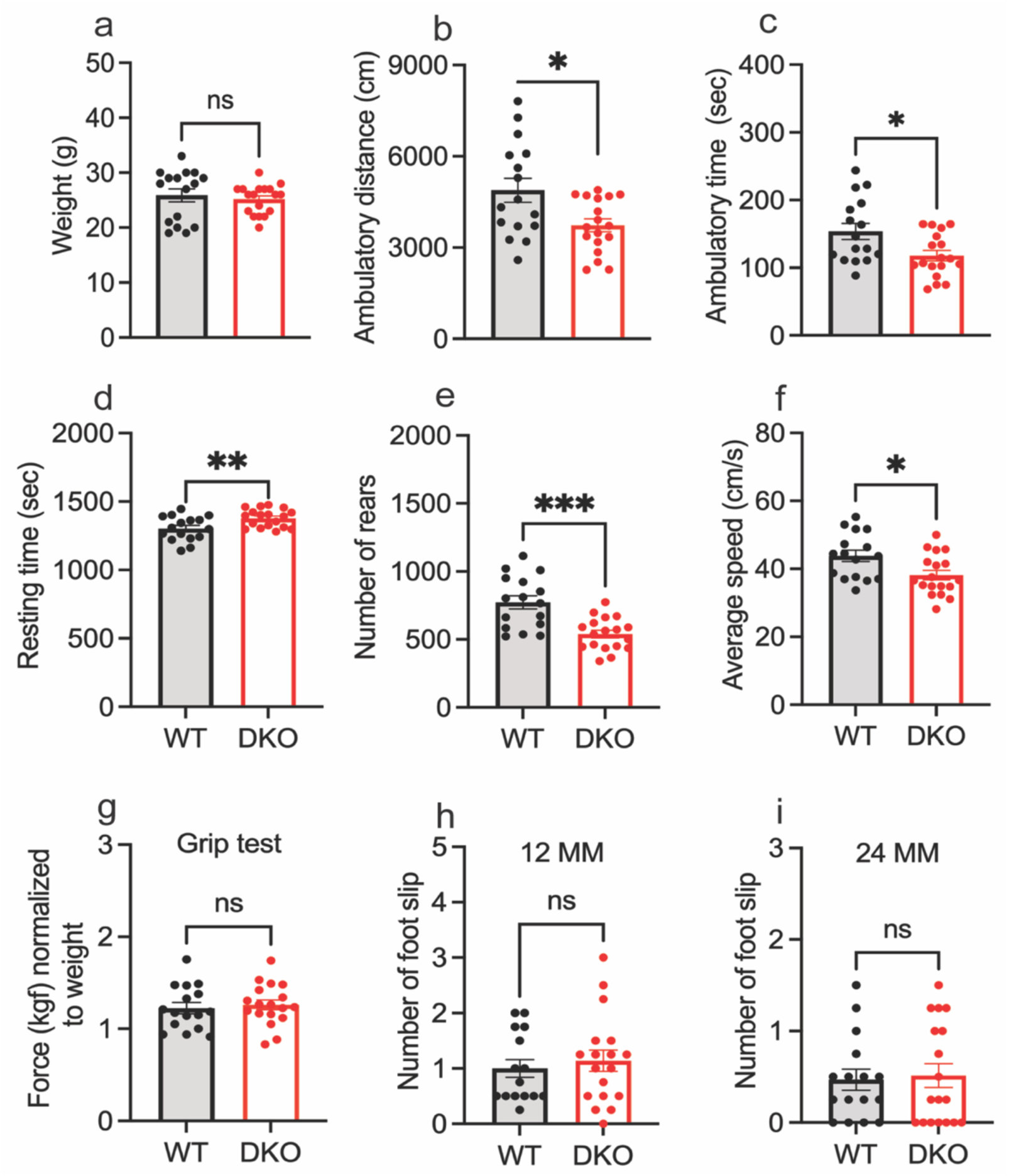
*Dnajc12* ablation impairs locomotion and exploratory behavior in the automated open field in 3M DKO mice. **a**) The weight of WT and DKO mice is comparable (Unpaired *t*-test with Welch’s correction, t_22.74_ = 0.53, p = 0.60). DKO mice have a significant reduction in their ambulatory distance (**b**, Unpaired *t*-test with Welch’s correction, t_23.16_ = 2.58, p = 0.02), ambulatory time (**c,** Unpaired *t*-test, t_32_ = 2.62, p = 0.01), number of rears (**e,** Unpaired *t*-test, t_32_ = 4.29, p = 0.0002) and average speed (**f**, Unpaired *t*-test, t_32_ = 2.58, p = 0.01), along with increased resting time (**d,** Unpaired *t*-test, t_32_ = 2.78, p = 0.001), all of which are indicative of impaired locomotion/exploratory behavior. Grip strength (**g,** Unpaired *t*-test, t_32_ = 0.46, p = 0.65) and sensorineural balance and coordination were comparable between WT and DKO (**h,** Unpaired *t*-test, t_32_ = 0.26, p = 0.80; **i**, Unpaired *t*-test, t_31_ = 0.54, p = 0.59). *\*p < 0.05, **p < 0.01, ***p < 0.001.* Error bars are ± SEM. 3M old mice, WT n = 16, DKO n = 18.

### Th protein and its phosphorylated forms are altered in MB and STR of DKO mice

As DNAJC12 is a chaperone for TH,^33^ we examined the consequence of its loss on Th protein and its phosphorylated forms, and additionally on the levels of heat shock cognate 70 (Hsc70) and 14-3-3 in MB and STR. Hsc70 may influence DA synthesis given it potentiates interactions with DA synthesizing enzymes, TH and aromatic amino acid decarboxylase (AADC), and with the vesicular monoamine transporter (VMAT2) which packages DA into vesicles.^34,35^ Similarly, 14-3-3 can modulate TH phosphorylation to control its activity, influencing DA synthesis.^36^ The protein expression of Hsc70 and 14-3-3 in MB and STR (Fig 4b-c, h-i) was comparable between DKO and their WT littermates, as were Th levels in STR (Fig 4d). However, in DKO STR, Th pSer31 was reduced while pSer40 was significantly increased compared to WT mice (Fig 4e-f). In contrast, total Th, pSer31 and pSer40 were all elevated in MB (Fig 4j-l).

**Fig 4.**
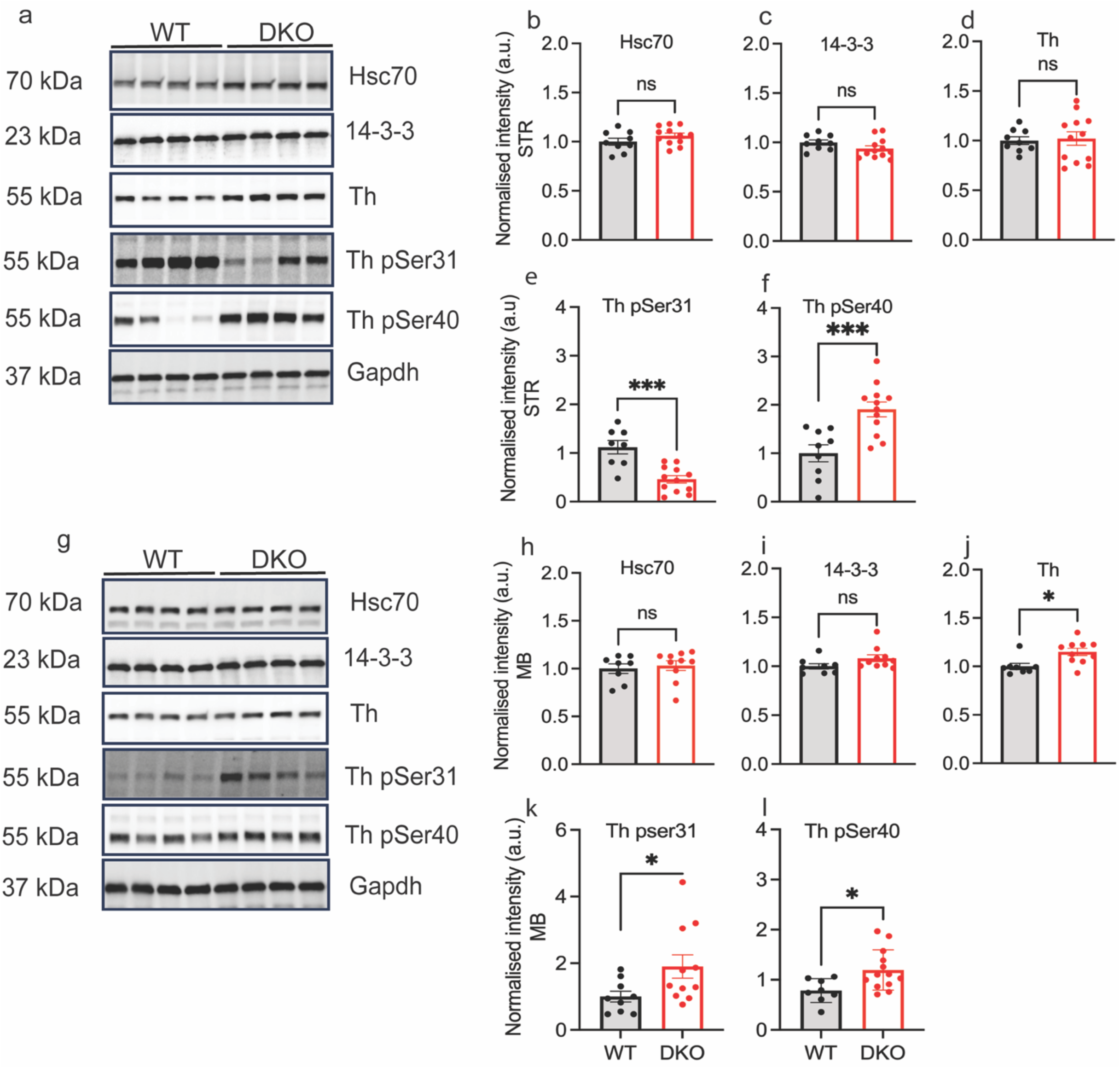
3M DKO mice exhibit alterations in Th protein, and its phosphorylated species in midbrain (MB) and striatum (STR). Representative blots for the STR and MB (**a, g**). The expression of Hsc70 (**b**, Unpaired *t*-test, t_19_ = 1.41, p = 0.17), 14-3-3 (**c,** Unpaired *t*-test, t_19_ = 1.57, p = 0.13) and Th (**d**, Unpaired *t*-test with Welch’s correction, t_16.68_ = 0.26, p = 0.79) in the striatum was comparable between WT and DKO mice. In contrast, pSer31 (**e**, Unpaired *t*-test, t_18_ = 4.52, p = 0.0003) was significantly reduced while pSer40 was increased (**f**, Unpaired *t*-test, t_19_ = 3.93, p = 0.0009). The expression of Hsc70 (**h**, Unpaired *t*-test, t_16_ = 0.43, p =0.67) and 14-3-3 (**i**, Unpaired *t*-test, t_16_ = 1.76, p = 0.09) was not altered in the midbrain in WT compared to DKO, meanwhile tTh (**j**, Unpaired *t*-test, t_16_ = 2.64, p = 0.01), Th pSer31 (**k**, Unpaired *t*-test with Welch’s correction, t_14_ = 2.332, p = 0.04) and Th pSer40 (**l**, Unpaired *t*-test, t_18_ = 2.70, p = 0.01) were significantly increased. *\*p < 0.05, ***p < 0.001.* Error bars are ± SEM. 3M old mice, WT n= 8-9, DKO n = 10-12.

### DKO mice have reduced striatal DA and electrically evoked peak response

Biogenic amines in micro-dissected STR were measured by HPLC. Striatal DA (43.70 versus 58.86 ng/mg), and its metabolites 3,4-dihydroxyphenylacetic acid (DOPAC) (3.20 versus 5.62 ng/mg) and homovanillic acid (HVA) (4.052 versus 6.823 ng/mg) were markedly reduced in DKO mice compared to their WT littermates (Fig 5b-e). DKO mice also exhibit reduced levels of striatal 5-HT and its metabolite 5-hydroxyindoleacetic acid (5-HIAA) (Fig S3b-d) and increased plasma levels of phenylalanine (Phe) (Fig S4a). However, other amino acids, such as tyrosine (Tyr) and tryptophan (Trp) (Fig S4 b-c), were unchanged. Reduced striatal 5-HT coincides with the role of Dnajc12 as a co-chaperone for TPH2, the rate-limiting enzyme in the synthesis of 5-HT in the central nervous system (CNS), while increased Phe levels denote the role of Dnajc12 as a co-chaperone for PAH, which catalyzes the conversion of Phe into tyrosine, an essential amino acid required for DA synthesis. Interestingly, loss of Dnajc12 didn’t change the levels of plasma biogenic amines as measured with mass-spectrometry (Fig S4d-g). To examine dynamic changes in striatal DA, we measured DA release and reuptake in *ex-vivo* striatal slices with FSCV (Fig 6a-c). When electrically stimulated, slices from DKO mice have a significantly lower peak response compared to WT mice (0.14 versus 0.30 μM) (Fig 6d), which concurs with reduced striatal DA. The decay constant tau was comparable between DKO and WT mice (415 versus 425 ms) (Fig 6e) signifying unaltered reuptake kinetics of DA by terminals.

**Fig 5.**
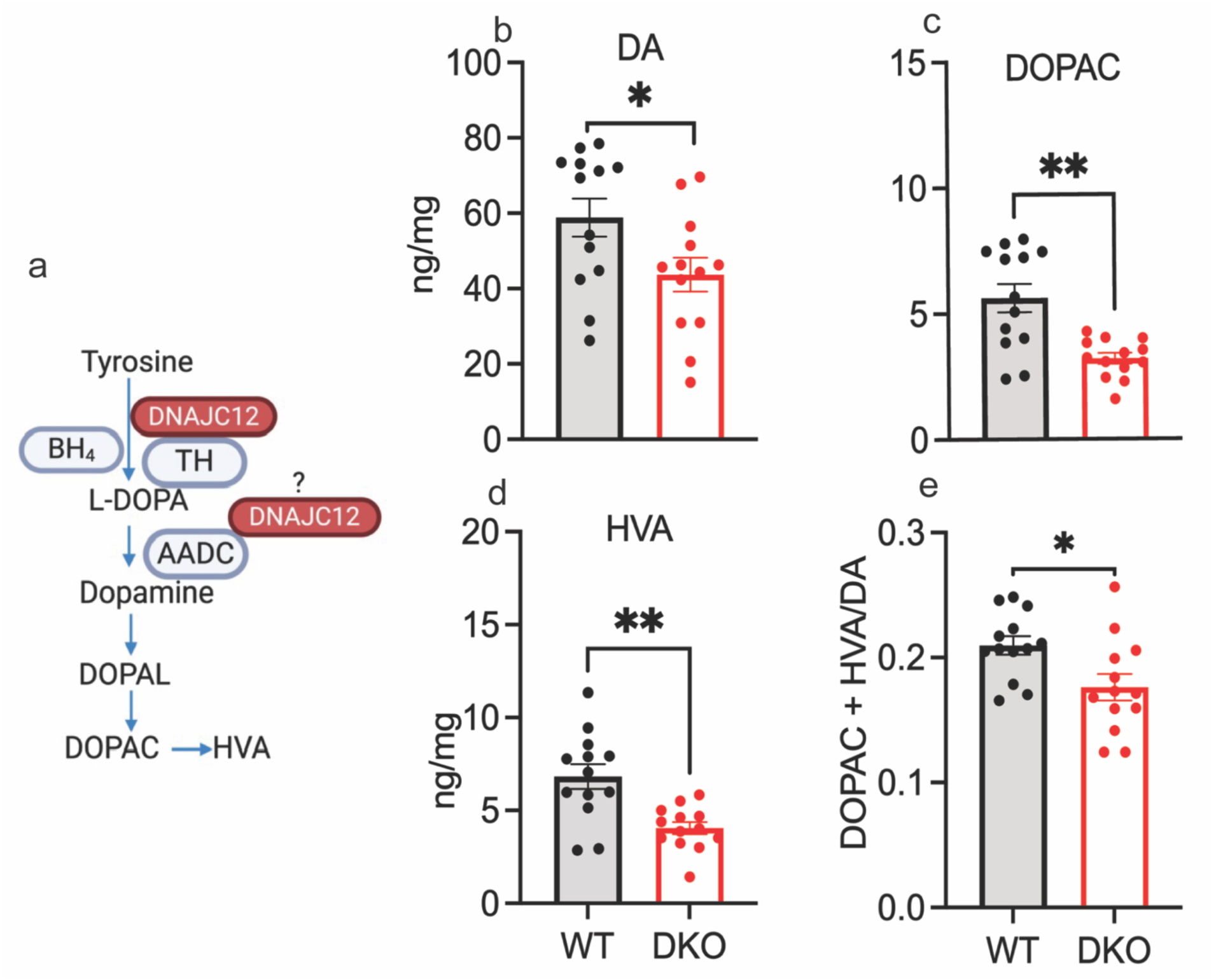
*Dnajc12* knock-out reduces the levels of striatal DA, and its metabolites in 3M DKO mice. **a**) Depiction of DA metabolism. Levels of DA (**b,** Unpaired *t*-test, t_24_ = 2.25, p =0.03), DOPAC (**c,** Unpaired *t*-test with Welch’s correction, t_15.43_ = 4.01, p = 0.001), HVA (**d,** Unpaired *t*-test with Welch’s correction, t_17.36_ = 3.74, p = 0.002) and ratio of DA metabolites over total DA (DOPAC + HVA/DA) (**e**, Unpaired *t*-test, t_24_ = 2.59, p = 0.016) via HPLC. *\*p < 0.05, **p < 0.01.* Error bars are ± SEM. WT n = 13, DKO n = 13.

**Fig 6.**
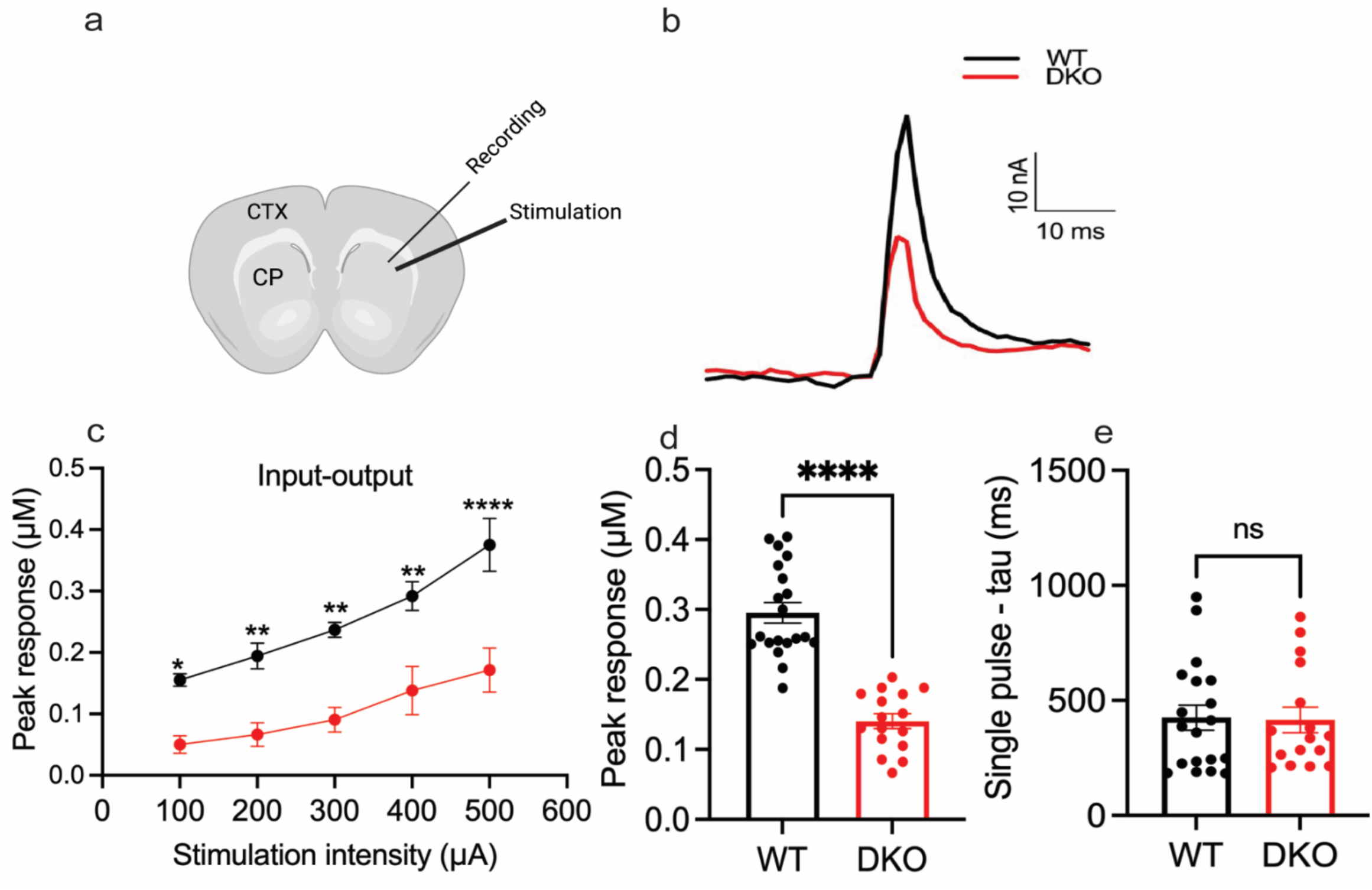
The peak amplitude of striatal DA release is reduced in 3M DKO mice. **e**) Depiction of the electrode’s placement into the striatal slices, n refers to the number of mice and in the parentheses the number of slices (WT n = 5 (20); DKO n = 4 (16)). Representative DA traces for WT and DKO (**f**). A series of increasing intensity single pulse stimuli were delivered every 2 min, and the input-output peak response is as indicated (**g,** starting from 100 to 500 µA, 2-way ANOVA with Šídák’s multiple comparisons test: t_29_ = 4.14, p = 0.001; t_29_ = 3.54, p = 0.007; t_29_ = 3.62, p = 0.006; t_29_ = 4.10, p = 0.002; t_29_ = 5.43, p = <0.0001). Single pulse peak response (stimulation at 400 µA) with 70% of the maximum DA release (**h,** Unpaired *t*-test, t_34_ = 8.23, p <0.0001). Single pulse tau (ms) (**i**, Unpaired *t*-test, t_33_ = 0.13, p = 0.90). *\*p < 0.05, **p < 0.01, ****p < 0.0001.* Error bars are ± SEM.

### Expression of endocytic and/or synaptic proteins is altered in STR of 3M DKO mice

DNAJC12 interacts with TH, VMAT2, AADC and HSC70, making it central to DA synthesis.^34,35^ As such, it’s loss could also alter endocytic proteins that foster the formation and recycling of synaptic vesicles, along with proteins involved in DA neurotransmission. Thus, we examined the expression of striatal endocytic and synaptic proteins including other DNAJC family members implicated in parkinsonism. There were no changes in proteins such as clathrin heavy chain (Chc) (Fig 7b), vesicular associated membrane protein 2 (Vamp2) (Fig 7e), and cysteine string protein-α (Dnajc5) (Fig 7f). In contrast, synaptosomal associated protein 25 (Snap25) (Fig 7d) was significantly reduced while endophilin 1A (Fig 7c) was increased in DKO mice compared to WT, indicative of major changes in axonal endocytosis and synaptic vesicle recycling.

**Fig 7.**
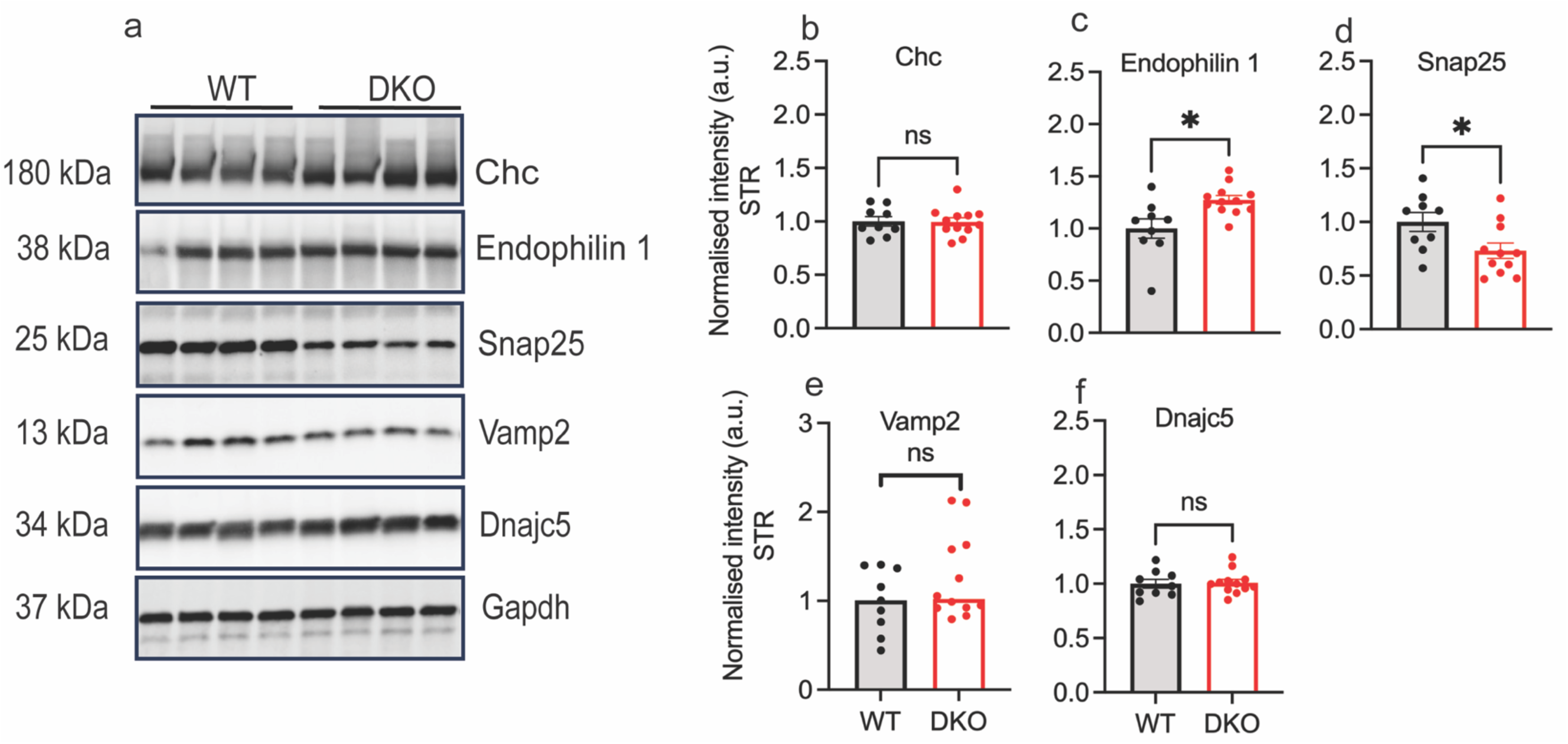
*Dnajc12* knock-out on the expression of synaptic proteins in the striatum (STR). Representative blots for the expression of synaptic proteins in the striatum (STR) (**a**). Densitometric analyses for clathrin heavy chain (Chc) (**b**, *Unpaired t*-test, t_19_ = 0.07, p = 0.94), endophilin 1 (**c**, Unpaired *t*-test with Welch’s correction, t_11.20_ = 2.70, p = 0.02), synaptosomal associated protein 25 (Snap-25) (**d**, *Unpaired t*-test, t_18_ = 2.35, p = 0.03), vesicular associated membrane protein 2 (Vamp2) (**e**, *Unpaired t*-test, t_19_ = 1.34, p = 0.19) and cysteine string protein-α (Dnajc5) (**f**, *Unpaired t*-test,t_19_ = 0.20, p = 0.84). *\*p < 0.05.* Error bars are ± SEM. 3M old mice, WT n= 9, DKO n = 12).

## Discussion

DA is critical in regulating movement, attention, addiction, reward, arousal, wakefulness and cognition.^37–40^ These functions are under the control of dopaminergic circuits, which include the nigrostriatal, mesocortical and mesolimbic pathways.^37^DA deficiency is a core feature of patients with *DNAJC12* mutations, congruent with this protein’s role as a co-chaperone for the DA synthesizing enzyme TH.^3–5^ Hence, deficits in the DA system are likely responsible for many (but not all) of the neurological problems observed in patients with *DNAJC12* mutations.^5,6,8,41,42^ Earlier studies with unbiased mass-spectrometry in HEK293 cells overexpressing DNAJC12 revealed its interaction with TH and TPH, and has been confirmed *in-vitro* by co-immunoprecipitation of transiently co-transfected GFP-tagged DNAJC12 with FLAG-TH or FLAG-TPH1 in HEK293 cells.^4,43,44^ We also confirmed that DNAJC12 co-immunoprecipitates Hsc70, consistent with prior immunoaffinity-mass spectrometry^43^, that may influence DA biosynthesis and packaging by its interaction with VMAT2, TH and AADC.^34,35^ We have also shown DNAJC12 interacts with GCH1, forming a complex with TH, and that this complex is destabilized when DNAJC12 is knocked-down, which markedly reduces levels of GCH1 without altering TH. Overexpression of both TH and GCH1 enhances levels of endogenous DNAJC12, suggesting it is required for the stability of the former enzymes/complexes. Hence, modulation of DNAJC12-GCH1-TH levels with pharmaceutical agents, such as sapropterin dihydrochloride to supplement BH_4_, or replacement of *DNAJC12* with adeno-associated viral delivery, may be of therapeutic benefit.

We extended *in*-*vitro* studies to assess the loss of Dnajc12 on DA biology, physiology and behavior in 3M DKO mice homozygous for the Dnajc12 p. V27Wfs*44 allele. This genetic modification is analogous to *DNAJC12* c.79-2A>G (p. V27Wfs*14), first discovered in a patient with young-onset parkinsonism and results in nonsense-mediated decay.^5^ Comparative analyses in 3M DKO and WT littermates indicate DKO have a significant reduction in locomotion and exploratory behavior. Bradykinesia is a core features required for a diagnosis of PD. However, motor deficits in patients with pathogenic *DNAJC12* variants are often preceded by neurodevelopmental and neuropsychiatric features, which may also explain reduced exploratory/locomotive behaviour in the DKO mice. Additional testing in 3M DKO mice, using an elevated-plus maze for anxiety-like behavior or Bussey Saksida boxes for cognition, might reveal additional behavioural phenotypes.

In PD, the degeneration of DA neurons in the SNpc appears to start in striatal terminals, and is accompanied by reduced TH protein/activity in remaining neurons.^45–47^ Nevertheless, the initial description of a patient with homozygous *DNAJC12* c.79-2A>G (p. V27Wfs*14) suggests an enzymatic form of parkinsonism for which longitudinal imaging revealed little evidence of progressive dopaminergic deficits.^5^ In DKO versus WT mice, striatal DA, its metabolites DOPAC/ HVA, and the amplitude of DA evoked release in *ex-vivo* slices were all significantly reduced. Thus we investigated the effects of Dnajc12 ablation on nigrostriatal DA synthesis by quantifying Th levels and phosphorylation in micro-dissected MB, and STR. While total Th, pSer31 and pSer40 were all elevated in the MB, in the STR in DKO compared to WT, pSer31 was reduced and only pSer40 was increased. Th phosphorylation at these serine residues correlates with activity but it was remarkable to see regional differences in terminals versus soma.^48,49^ In DKO mice, the loss of Dnajc12 appears to alter the interaction, stability and activity of Th, and culminates in reduced DA levels and behavioral deficits. Notably, increased Th phosphorylation in MB and STR of DKO mice may reflect compensation to modulate its activity to meet the demands of DA synthesis from the loss of Dnajc12. As a corollary, striatal synaptic proteins such SNAP25 which directly affects vesicle exocytosis, SNARE complex formation and neurotransmission^50–52^, and endophilin 1 which influences vesicle endocytosis, mitochondrial morphology and autophagy^53–57^, are also altered in DKO mice and potentially reflect compensation within and beyond DA terminals. It remains to be seen if DA terminal and axonal degeneration will occur in aged DKO mice.

DKO mice also have reduced levels of striatal 5-HT, attributed to the role of DNAJC12 as a co-chaperone for TPH2, the rate-limiting enzyme for the synthesis of 5-HT in the CNS and the periphery.^5,58^ 5-HT in the CNS is critical for neurite outgrowth, synaptogenesis and neurogenesis.^59,60^ In contrast, peripheral 5-HT has a role in gastrointestinal motility and inflammation^61–64^, while adrenal catecholamines (including DA, adrenaline and noradrenaline) are essential in autonomic responses, secreted into the bloodstream as hormones with effects on many peripheral organs.^37,65–67^ Nevertheless, changes in peripheral biogenic amines may need be evoked with stress stimuli (fight or flight) as plasma biogenic amines were comparable between DKO and WT only. Mild HPA is a cardinal feature in human patients with pathogenic *DNAJC12* variants, and DKO mice have increased levels of plasma phenylalanine.

Dystonia and PD are considered phenotypically, physiologically and molecularly distinct but the etiologies in patients with *DNAJC12*^3–5,21,22^ or *GCH1*^15,17,23,24^ mutations overlap, and many features can be attributed to DA dysregulation, as both are responsive to L-DOPA treatment. Enzymatic pathways of catecholamine synthesis are well documented, but the composition and relationship of many molecular components is unknown. Here we characterize DKO mice, which manifest reduced locomotion/exploratory behavior and biogenic amines levels, and mild HPA, which recapitulates the effects of *DNAJC12* mutations in human disease.^9^ Nevertheless, as constitutive *Dnajc12* loss may perturb the synthesis of other biogenic amines, we also developed homozygous cDKO mice to enable tissue and cell-specific ablation of Dnajc12 with somatic Cre recombinase. For example, Dnajc12 has been successfully ablated in MB and STR via bilateral intrastriatal injection of *AAV_2_ TH: Cre-eGFP* in cDKO mice (Fig S2a-e). Hence, DKO mice and their floxed founders, cDKO, provide preclinical models to study this pathobiology, and to test therapeutic strategies to increase TH protein/activity or DA synthesis. Interestingly, studies in PD report reduced BH_4_ concentration and GCH1 activity in the *SNpc* and STR, and low BH_4_ levels in cerebrospinal fluid (CSF).^45,68^ BH_4_ may also modulate TH levels and activity but nevertheless, preclinical experiments have yet to prove conclusive.^45,68^ Increased levels of BH_4_ may stimulate TH activity and DA release^69–71^, or might be toxic to catecholamine neurons^72,73^ Hence, DKO/cDKO mice will prove a valuable tool to uncover the mechanism, timing and therapeutic effects of sapropterin dihydrochloride supplementation (BH_4_) *in-vivo.* Despite many more patients with DNAJC12-deficient HPA being identified through the addition of *DNAJC12* genetic testing in newborn, the optimal treatment for patients has yet to be defined and remains an unmet need.^74^

## Supplementary figure legends

**Fig S1.**
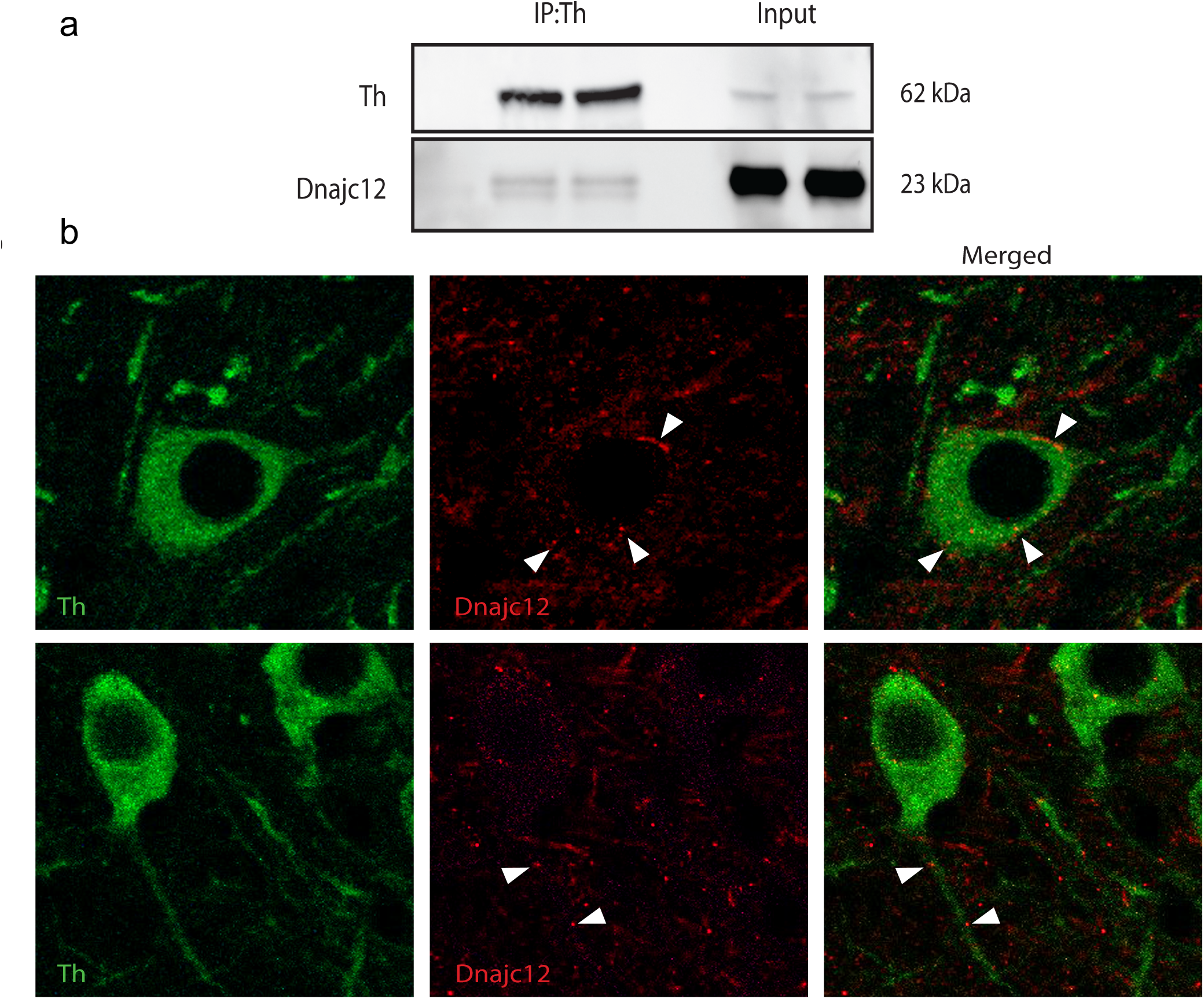
Dnajc12 which interacts with Th *in-vivo* localizes to the soma, dendrites and axons in Th-positive neurons. **a)** Co-immunoprecipitation assay depicting Dnajc12 interaction with Th. **b**) Immunofluorescent labelling with confocal microscopy showing Dnajc12 puncta in the soma, dendrites and axons in Th-positive neurons in coronal slices obtained from WT C57BL/6 mice.

**Fig S2:**
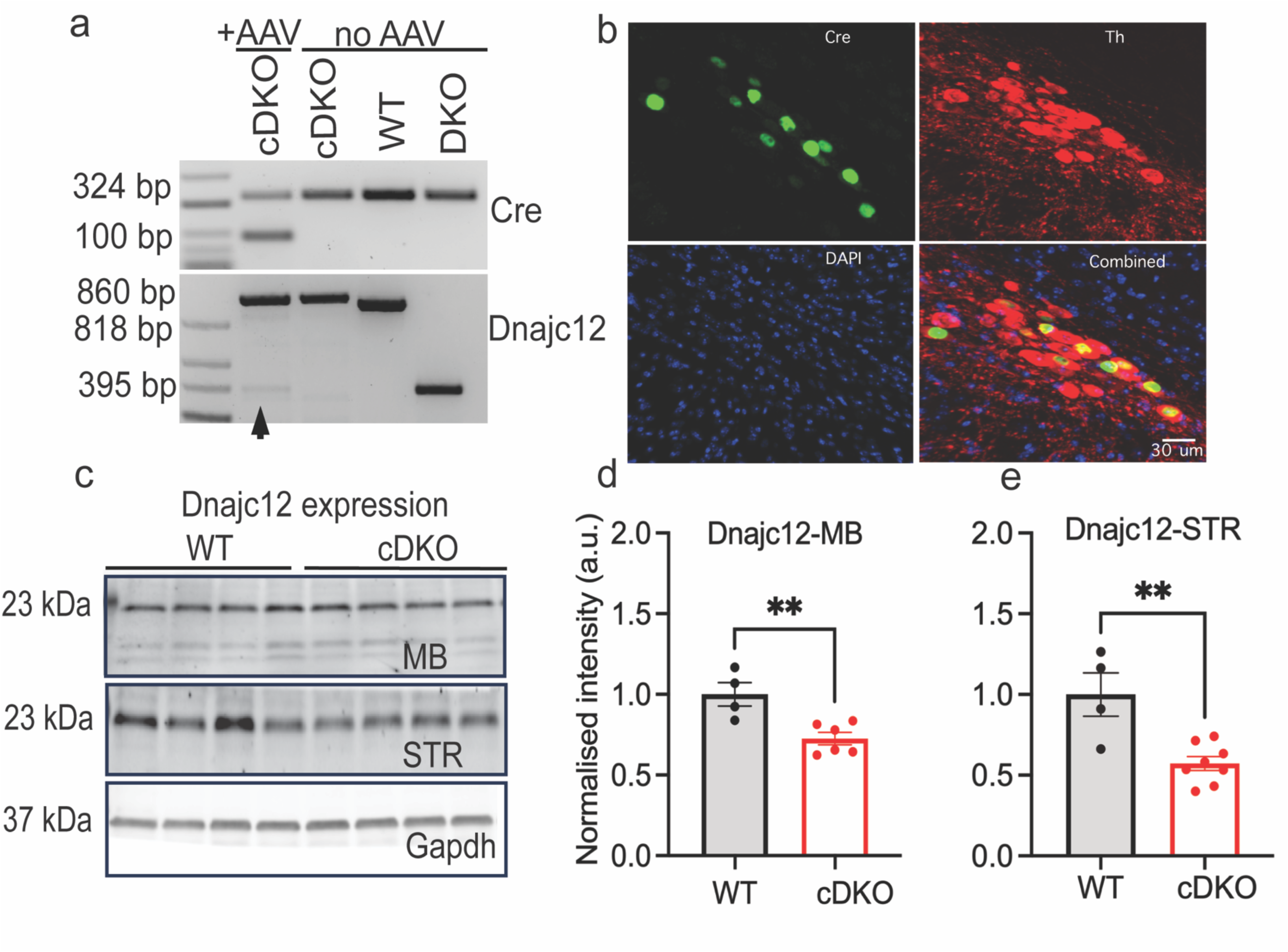
Ablation of *Dnajc12* in *SNpc* dopaminergic neurons and their projections to the striatum via intrastriatal injection of AAV_2_ TH: Cre-eGFP in homozygous 3M cDKO mice. a) Genotyping of gDNA control (300 bp), AAV_2_ Cre (100 bp), and *Dnajc12* for WT (818 bp), DKO (395 bp), WT (floxed) (860 bp) and cDKO mice (860 and 395 bp) in striatum. Representative immunoblots (**b**) and densitometric analyses of reduced Dnajc12 in MB (**c,** Unpaired *t*-test t_8_ = 3.67, p=0.007) and STR (**d,** Unpaired *t*-test, t_10_ = 3.89, p = 0.003). Error bars are ±SEM. WT n=4; cDKO n=7-8. Cre-expression in Th+ MB neurons from *AAV_2_ TH: Cre-eGFP (e)*, IF with confocal microscopy at 20x indicating Th+ neurons (red), Cre (green) and Dapi (nuclei stain, blue).

**Fig S3.**
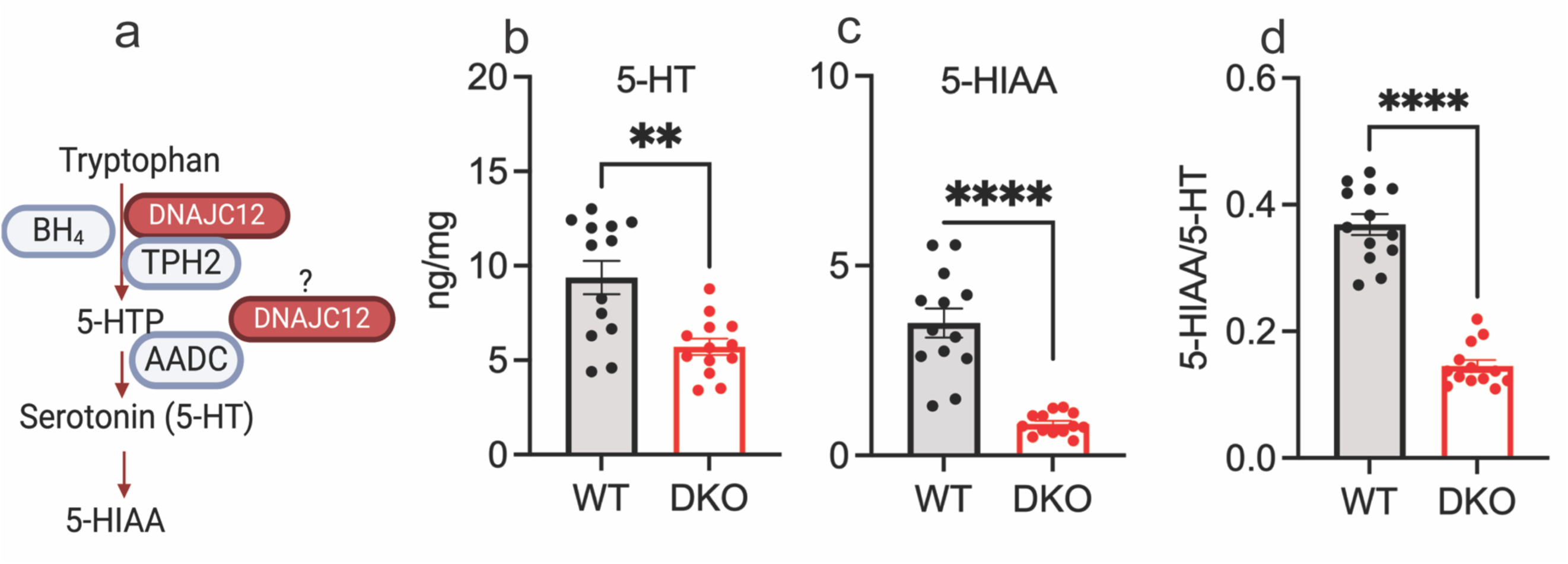
Striatal serotonin (5-HT) levels and its metabolite are reduced in 3M DKO mice. Depiction of serotonin synthesis pathway (**a**). Levels of 5-HT (**b**, Unpaired *t*-test with Welch’s correction, t_17.36_ = 3.74, p = 0.002), its metabolite 5-HIAA (**c**, Unpaired *t*-test with Welch’s correction, t_13.06_ = 6.85, p < 0.001) and ratio of 5-HT metabolite over total 5-HT (5-HIAA/5-HT) (**d,** Unpaired *t*-test, t_24_ = 11.79, p = *p < 0.0001)*. **p < 0.01, ***p < 0.001, *\*\*\*\*p < 0.0001*. Error bars are ±SEM. (WT n = 13; DKO n=13).

**Fig S4.**
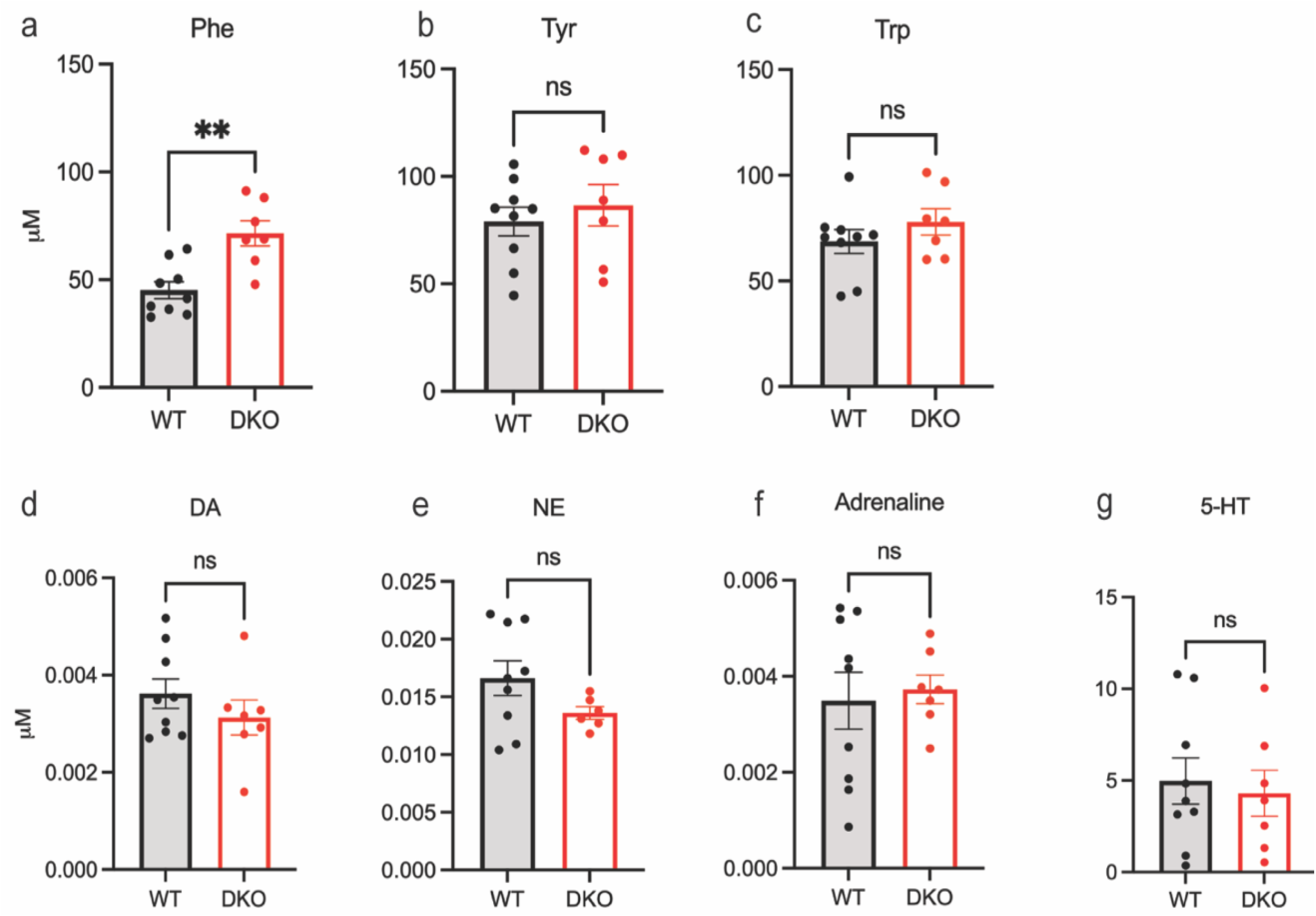
*Dnajc12* knock-out recapitulates hyperphenylalaninemia as exemplified in human patients with similar pathogenic variants but doesn’t alter plasma biogenic amines. Levels of plasma phenylalanine (Phe) (**a,** Mann-Whitney test, p = 0.005), tyrosine (Tyr) (**b,** Mann-Whitney test, p = 0.47), tryptophan (Trp) (Mann-Whitney test, p = 0.41) and plasma biogenic amines which include DA (**d**, Mann-Whitney test, p = 0.31), NE (Mann-Whitney test, p = 0.18), Adrenaline (**f**, Mann-Whitney test, p >0.10) and 5-HT (**g**, Mann-Whitney test, p = 0.84). **p < 0.01. Error bars are ±SEM. 6-8M old mice, WT n=9, DKO n=7.

**Fig S5.**
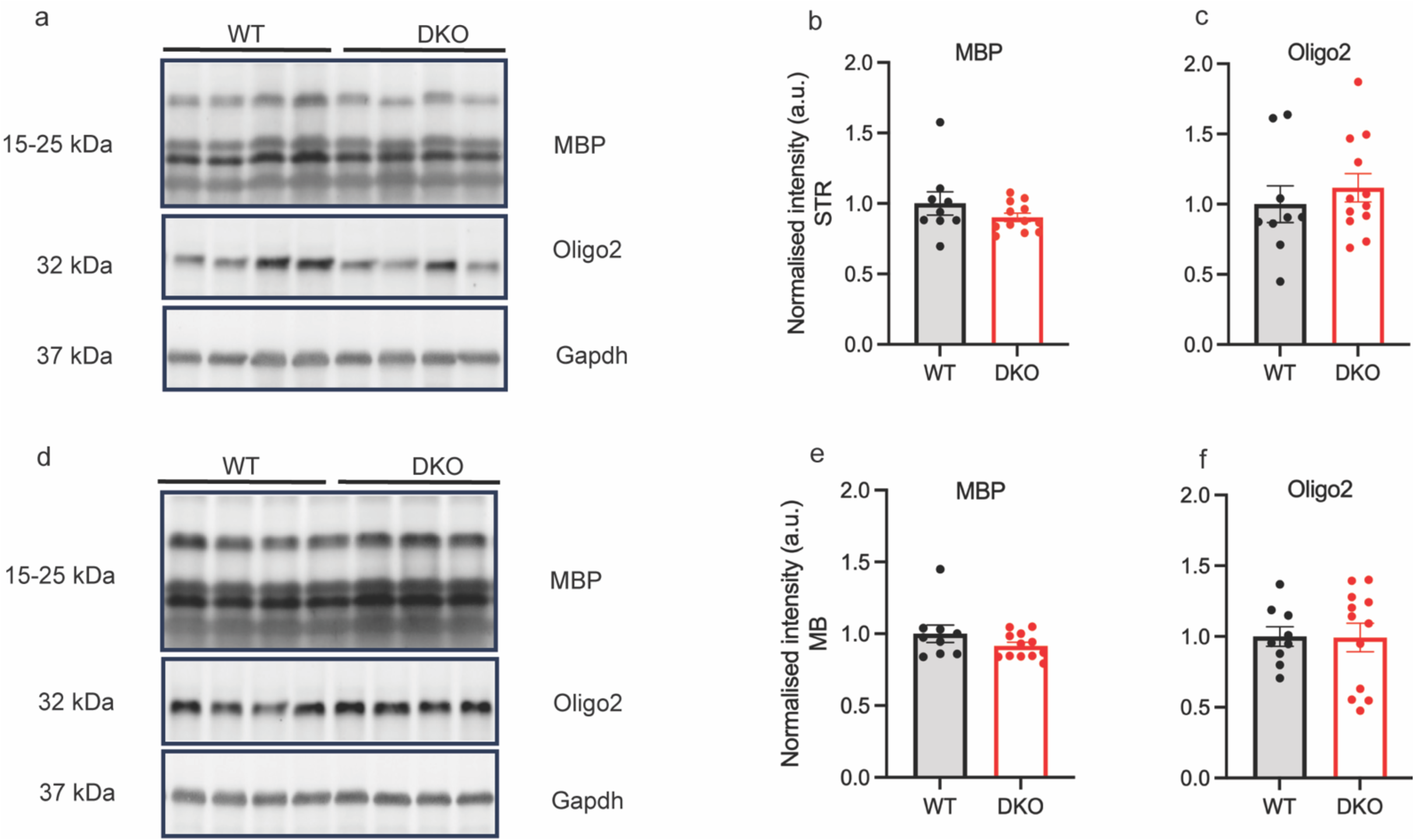
Oligodendroglial markers are unchanged in the striatum (STR) and midbrain (MB) of 3M DKO mice. Representative blots for MBP and Olig2 in the STR (**a)** and MB (**d**). Densitometric analysis of MBP-STR (**b**, Unpaired *t*-test with Welch’s correction, t_10.16_ = 1.14, p = 0.28), Olig2 – STR (**c**, unpaired *t*-test, t_19_ = 0.72, p = 0.48), MBP-MB (**e**, Unpaired *t*-test with Welch’s correction, t_10.75_ = 1.27, p = 0.23) and Olig2-MB (**f**, unpaired *t*-test, t_19_ = 0.06, p = 0.96). Error bars are ±SEM. (WT n = 9; DKO n=12).

## Acknowledgement

This study was supported in part by the Emory HPLC Bioanalytical Core (EHBC), which was supported by Emory University School of Medicine and the Georgia Clinical & Translational Science Alliance of the National Institutes of Health under Award Number UL1TR002378.

## Author contribution

IBD: conceptualization and project design, data curation, formal analysis, writing original draft, review and editing; JF: conceptualization and project design, data curation and formal analysis, review and editing; JDF: conceptualization and project design, data curation and formal analysis, MJF: Resources, conceptualization and project design, training and supervision, critical review, interpretation and editing.

## Notes

### Competing Interest Statement

The authors have declared no competing interest.

### Summary of Updates

– An additional author. – The text has been rewritten immensely to improve clarity and to enhance knowledge of the topic of interest. – New results that were not in the previous manuscript e.g Fig 1, 5, 6, Fig S2 and 3.

